# Ordered, Random, Monotonic, and Non-monotonic Digital Nanodot Gradients

**DOI:** 10.1101/001305

**Authors:** Grant Ongo, Sébastien G. Ricoult, Timothy E. Kennedy, David Juncker

**Affiliations:** Department of Biomedical Engineering, McGill University, 740 Dr. Penfield Avenue, Montreal, Quebec, Canada H3A 0G1; McGill University & Genome Quebec Innovation Centre, McGill University, Montréal, Quebec, Canada H3A 0G1; McGill Program in Neuroengineering, Department of Neurology and Neurosurgery, Montreal Neurological Institute, McGill University, 3801 University Ave, Montreal, Quebec, Canada H3A 2B4

## Abstract

Cell navigation is directed by inhomogeneous distributions of extracellular cues. It is well known that noise plays a key role in biology and is present in naturally occurring gradients at the micro- and nanoscale, yet it has not been studied with gradients *in vitro*. Here, we introduce novel algorithms to produce ordered and random gradients of discrete nanodots – called digital nanodot gradients (DNGs) – according to monotonic and non-monotonic density functions. The algorithms generate continuous DNGs, with dot spacing changing in two dimensions along the gradient direction according to arbitrary mathematical functions, with densities ranging from 0.02% to 44.44%. The random gradient algorithm compensates for random nanodot overlap, and the randomness and spatial homogeneity of the DNGs were confirmed with Ripley’s K function. An array of 100 DNGs, each 400 × 400 µm^2^, comprising a total of 57 million 200 × 200 nm^2^ dots was designed and patterned into silicon using electron-beam lithography, then patterned as fluorescently labeled IgGs on glass using lift-off nanocontact printing. DNGs will facilitate the study of the effects of noise and randomness at the micro- and nanoscales on cell migration and growth.

## Introduction

Gradients are fundamental to many phenomena of biology, from directing axonal navigation during neural development to the differentiation of stem cells in response to an injury [1,2]. Gradients may occur as either (i) free diffusion gradients, (ii) substrate-bound gradients, or (iii) a combination of both [3]. Free diffusion gradients can be generated by cell-excreted proteins that diffuse in the extracellular space. Substrate-bound gradients arise when proteins are bound to the extracellular matrix or cell membranes [4,5,6]. To better understand cellular haptotaxis – the directed movement of a cell or axonal growth cone along a gradient of a substrate-bound guidance cue – many techniques to generate surface-bound gradients *in vitro* have been developed [7]. Two classes of concentration gradients exist, namely continuous and digital gradients. Continuous gradients have protein concentration changing in a constant manner, and are typically produced by adsorbing molecules from a diffusible gradient. Several parameters of these gradients have been modulated, including the range and slope, which was linear, exponential or non-monotonic [8]. For example, linear gradients of the protein BDNF have been shown to modulate neuron polarization and growth [9] and linear laminin gradients have been shown to orient rat hippocampal axon specification based on slope [10]. Secondly, digital gradients which have been recently introduced, are formed by patterning small dots of protein and varying the density of the dots by changing their size [11], the spacing between them [12], or both [13]. The advantage of digital gradients is that they are deterministic, and that the local concentration can be precisely predicted and does not rely on fluorescence measurements which are prone to error. Digital gradients have been used to study how retinal ganglion cells identify the stop zone within graded distributions of repulsive EphrinA5 ligands [13]. However, digital gradients rarely extend over 1-2 orders of magnitude (OM), whereas it is believed that the dynamic range of gradients *in vivo* is between 3-4 OM. To overcome these limitations, we previously developed digital nanodot gradients (DNGs), where the spacing between nanodots (200 nm in diameter) was changed in two dimensions to produce a dynamic range exceeding 3 OM. These designs were implemented using a low-cost, lift-off nanocontact printing method to pattern substrate-bound gradients of proteins and peptides. We employed these patterns for adhesion and migration studies of C2C12 myoblasts on RGD peptide and netrin-1 gradients, respectively [14]. For these experiments, gradients of 400 × 400 µm^2^ were divided into 53 rectangular boxes of fixed size and density. With this approach, gradients were non-continuous and had stepwise changes in density. This is most pronounced in low-density regions where the spacing between nanodots exceeded the dimensions of the box, requiring larger box sizes and thus creating large “steps” at the bottom of the gradient. This may prove problematic as cells may fail to sense a discontinuous gradient if they fall into a region of constant density.

Noise is ubiquitous in biology [15], and modern patterning technologies can be exploited to introduce defined amounts of noise and randomness into otherwise regular patterns. The effect of randomness was evaluated in ordered arrays and repetitive patterns with a constant average density. In one study, disorder was introduced in arrays of 120 nm dots spaced 300 nm apart by randomly displacing the dot by up to 50% of the spacing to avoid overlap [16]. While this approach introduced a controlled amount of noise by restricting the maximum displacement, it was not random since dots were each contained within the original grid. Nonetheless, cellular adhesion and stem cell differentiation of osteoblasts were markedly altered as a result of increasing disorder. In another study, whole proteome analysis of cells grown on the same disordered patterns resulted in differential expression of certain proteins in the extracellular signal-regulated kinase (ERK1/2) pathway [17]. Similarly, controlled amounts of topographical noise on nanogratings of 500 nm ridges and grooves has shown to effect PC12 neuronal cell alignment to the gratings, focal adhesion maturation and directionality [18].

Randomness and noise are also highly relevant to directed cell migration. The stochasticity of chemo- and haptotaxis has been well studied, and is apparent from the random-walk like traces of migrating cells [19]. It is well understood that biological gradients, which appear continuous, are in fact quantized since they are comprised of individual molecules adsorbed to a surface. The distribution of these molecules is not deterministic, but stochastic at the nanoscale. The engagement of receptors from migrating cells with these guidance cues has been modeled within a stochastic framework [19,20]. Random variations in the gradient also occur at the microscale *in vivo* from the local accumulation of chemo- and haptotactic molecules that form concentration puncta [21]. Gradients formed by the expression of receptors from cells embedded in a tissue a may become non-monotonic as particular cells over- and under-express a receptor relative to their position in the gradient [22]. Cells navigating through such a patchwork of microscale deviations must discriminate against local maxima and minima to sense the overarching gradient slope. It has been suggested that cells alternate between periods of sensitization and desensitization and are thereby capable of maintaining an overall response to long-range gradients [23].

Random pattern generation is well established, however many patterns are not random in the mathematical sense, and often implement only an approximation of randomness to varying degrees depending on the needs of the application. Firstly, computers typically generate ‘pseudo-random’ numbers that statistically approximate the properties of true random numbers using an algorithm based on an initial ‘random seed’ – generally a random bit. If this random seed is known, the set of pseudo-random numbers can be replicated. However, for most applications, pseudo-random numbers are indistinguishable from a true random number.

So-called random dot patterns are widely used in the reproduction of images and patterns using displays or printers that impose digitization and pixilation. Pixilation implies that pixels are positioned on a regular grid. Adjusting grayscale images to binary (black and white pixels) through halftoning is achieved by adjusting the ratio of black and white pixels in a local area, but if pixels are turned on and off according an ordered pattern, rendering artifacts (such as Moiré patterns) arise. By locally randomizing the pixels turned on or off while matching the overall desired grayscale, it is possible to avoid such rendering artifacts. These ‘random-grid’ patterns can be generated using an array of pseudo-random values and a threshold dictated by the local grey value used to determine which pixels are black or white (Fig. S1) [24]. With this approach, the value of each pixel is random, while the position of the pixels are fixed to an ordered grid. These random-grids have found application in the halftoning of images for display or printing [25] and the generation of random-dot stereograms for depth perception studies [26].

Random dot patterns also comprise dot patterns where the position of the dots is not constrained to a grid as described previously, but is instead randomized within a given space. However, pseudo-random distributions of the dots, when seen from afar, can appear noisy and inhomogeneous to the eye. Therefore, algorithms involving so-called quasi random numbers (also called low discrepancy sequences) are used to create patterns with a more uniform distribution that appears visually smooth [27]. “Quasi-random” is a broad term that encompasses numbers with a disordered distribution that lie between a (pseudo-)random distribution and a regular distribution [28]; a quasi-random distribution may closely resemble an ordered distribution than a random distribution upon close examination. Many different algorithms have been developed to produce quasi-random numbers. As an example, for liquid crystal displays, quasi-random dot patterns without overlap are generated by mimicking molecular dynamics in which two “molecules” cannot occupy the same space [29] or by iteratively repositioning overlapping dots until overlap is minimized [27].

Both random-grid and quasi-random patterns have been optimized to produce a pleasing (macroscopic) image to the eye by introducing a controlled amount of microscale disorder, avoiding both the viewing artifacts of regular arrays and the inhomogeneity of pseudo-random dot patterns. Cells however are microscopic and sense their environment at the micro- and nanoscale. At these scales, random grid or quasi-random distributions can be significantly different from a random one, and may not adequately reproduce the effect of a random pattern on cells. It is therefore necessary to establish an algorithm to produce random dot patterns that preserves randomness at the micro- and nanoscale to study the effect of random *vs.* regular dot patterns on cell navigation.

Here, we introduce continuous DNGs eliminating the stepwise density changes of the previously reported “step” design [14] with (i) ordered and (ii) pseudo-random positioning of nanodots, as well as (iii) non-monotonic DNGs that can be implemented using either (i) or (ii). We discuss various strategies to create noisy DNGs and outline the challenges in forming random DNGs with accurate slopes, and describe a novel algorithm with pseudo-random positioning of the dots and compensating for random, but predictable, dot overlap to achieve the desired coverage. The slope of random DNGs was measured and compared to the programmed density function, and their randomness verified using Ripley’s K-function. We generated an array of 100 ordered and random 400 × 400 µm^2^ large DNGs made of 200 × 200 nm^2^ nanodots, including monotonic and non-monotonic density curves, with a dynamic ranges spanning from 2.14 to 3.86 OM. Non-monotonic gradients produced here aim to introduce in a quantitative and repeatable manner “microscale noise” into surface-bound *in vitro* gradients. The entire array of 100 DNGs covers a 35 mm^2^ area and is comprised of more than 57 million nanodots. The DNG array was patterned onto a silicon (Si) wafer by electron-beam (e-beam) lithography, and was transferred onto glass slides by lift-off nanocontact printing of fluorescently labeled IgGs [14]. The fidelity of the replication process was evaluated by overlaying the DNG design with fluorescence microscopy images of the printed IgG proteins.

## Materials and Methods

### Software & Computers

Algorithms were developed in Matlab R2013a (Natick, MA, USA). Scripts of the ordered and random gradient algorithms are available for download upon request. A spreadsheet template (Microsoft Excel 2010) of gradient parameters was imported into Matlab for processing. The output for each gradient was a text file of coordinates formatted as a Caltech Intermediate Format (CIF) file. CIFs were imported into CleWin Version 4.1 (WieWeb, MESA Research Institute at the University of Twente and Deltamask, Netherlands). Gradients were exported from CleWin as Bitmap (BMP) image files for verification of density using ImageJ 1.47 64-bit (National Institutes of Health, USA) and Matlab. The 100-gradient array was designed in L-edit (Tanner EDA, Monrovia, CA, USA). All software was run on a 2013 iMac computer with a 3.4 GHz Intel Core i7 processor with 32 GB 1600 MHz DDR3 memory running Windows 7 through a bootcamp partition.

### Electron-Beam Lithography

A 4” Si wafer was coated with PMMA resist and the 100-gradient array was patterned by e-beam lithography (VB6 UHR EWF, Vistec, Montreal, QC, Canada), followed by reactive-ion etching (System100 ICP380, Plasmalab, Everett, WA, USA) 100 nm deep into the Si wafer.

### Stamp Fabrication

After cleaning, the Si wafer was coated with perfluorooctyltriethoxysilane (Sigma-Aldrich, Oakville, ON, Canada) by vapor phase deposition. An accurate polymer copy of the wafer was obtained after double replication using polydimethylsiloxane (PDMS) and a UV-sensitive polyurethane as described in [14]. First, a 6 mm layer of 1:10 PDMS (Dow Corning, Corning, NY, USA) was poured on the wafer inside a Petri dish, followed by degassing under vacuum in a desiccator for 10 min. Next, the PDMS was cured in an oven for 24 h at 60 °C (VWR, Montreal, QC, Canada), then peeled from the wafer. To remove uncured monomers and other extractables, the PDMS replica was submersed in 70% ethanol for 24 h then baked at 60°C for 4 h. Next, a drop of UV sensitive polyurethane (Norland Optical Adhesive 63 (NOA 63), Norland Products, Cranbury, NJ) was applied to the PDMS replica and cured by 600 W of UV light (Uvitron International, Inc., West Springfield, MA) for 30 s. The PDMS was then removed yielding an NOA replica of the original Si wafer pattern with 200 nm holes [14].

### Nanocontact Printing

A flat PDMS stamp was cured against a perfluorooctyltriethoxysilane treated flat Si wafer. Following removal of the extractables as mentioned above, the flat PDMS stamp was inked with a 10 µL drop of phosphate buffered saline solution (PBS) containing 25 µg/mL of chicken immunoglobulin G (IgG) conjugated to Alexa Fluor 488 (Invitrogen, Burlington, ON, Canada). A plasma activated hydrophilic coverslip was then placed on the drop to spread the solution evenly across the surface of the hydrophobic PDMS stamp during a 5 min incubation period. After rinsing with PBS and double distilled water for 15 s each, the inked stamps were briefly dried under a stream of N_2_ and immediately brought into contact with a plasma activated (PlasmaEtch PE-50, PlasmaEtch, Carson City, NV, USA) NOA master for 5 s. The PDMS and NOA were separated and proteins in the contact regions were transferred to the NOA. The remaining proteins on the PDMS were transferred to the final substrate by printing the PDMS stamp for 5 s onto a plasma activated glass coverslip.

### Imaging and Analysis

Images of the original Si master were collected using scanning electron microscopy (SEM, JEOL, Japan). DNGs of fluorescent IgGs were imaged by fluorescence microscopy (TE2000 microscope, Nikon, Canada and CoolSNAP HQ^2^ camera, Photometrics, USA). Images of the Si wafer were captured with a Panasonic Lumix GH3 DSLR equipped with an Olympus M. Zuiko Digital ED 60 mm macro lens. Dark field images were captured with an inspection microscope (LV150A microscope and Digital Sight DS-Fi1 camera, Nikon, Canada).

## Results and Discussion

### Ordered Gradient Algorithm

Ordered DNGs with continuously changing density were programmed by forming columns of nanodots with equal vertical spacing while varying the spacing between columns horizontally (Fig. 1A). The density of the gradient at any point along the length *l* is dictated by an input density function *D*, and is realized by placing a single nanodot into a virtual box to form a unit cell. The dimensions *d_i_* of the unit cell at the *i^th^* column of nanodots are given by the square root of the nanodot area *A_dot_* divided by the density value at the given position (Eq. 1). In this algorithm, the size of the nanodot remains constant while the dimensions of the unit cell vary for each column of nanodots along the length.

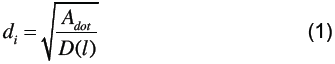

**Figure 1:**
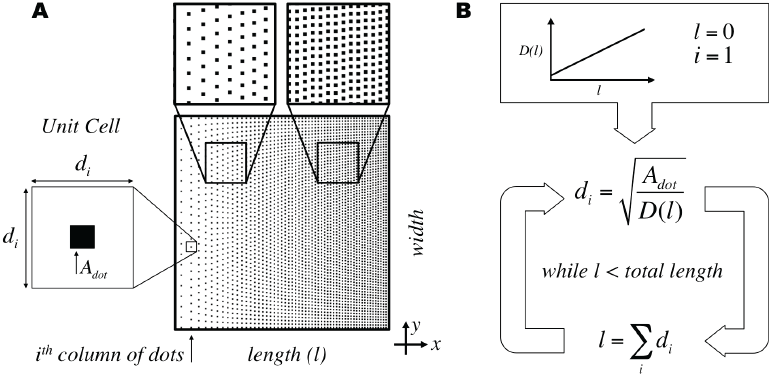
Ordered gradient algorithm for continuously changing density in Digital Nanodot Gradients (DNGs) using a unit cell approach. (A) Schematic of a linear DNG and unit cell parameters for the i^th^ column of nanodots. (B) Calculation of unit cell dimensions at each position through an iterative process using the cumulative sum of unit cell dimensions to determine position along the length.

The unit cell is largest at low densities and decreases at higher densities, matching the dimensions of the nanodot at a density equal to one. The unit cell dimensions for each column are calculated using an iterative approach, starting from the first column of nanodots at length zero. The position along the length for the next column of nanodots is given by the cumulative sum of unit cell dimensions (Fig. 1B).

The width of the gradient is divided by *d_i_* to estimate the number of nanodots in the column. The value of *d_i_* is then recalculated to equally space the integer number of nanodots along the width, ensuring that the distribution of nanodots is symmetrical. The unit cells are concatenated to form a column of points in the *y* direction with constant spacing as multiples of *d_i_*. The *x* coordinate for each column is the position *l* at which the unit cell dimensions are calculated. Thus, using this algorithm it is easy to form a gradient with any slope as the size of the unit cell is directly derived from the value of the density function at a specific position. Fig. 2 shows a linear density curve spanning from a minimum density *D_min_=0.01* to a maximum density *D_max_=0.30* (Eq. 2) with corresponding unit cell dimensions along the length of the gradient. Eq. 2 is normalized to span the dynamic range over the total length *L=100 µm* of the gradient.

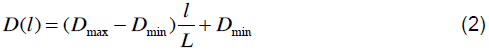

**Figure 2:**
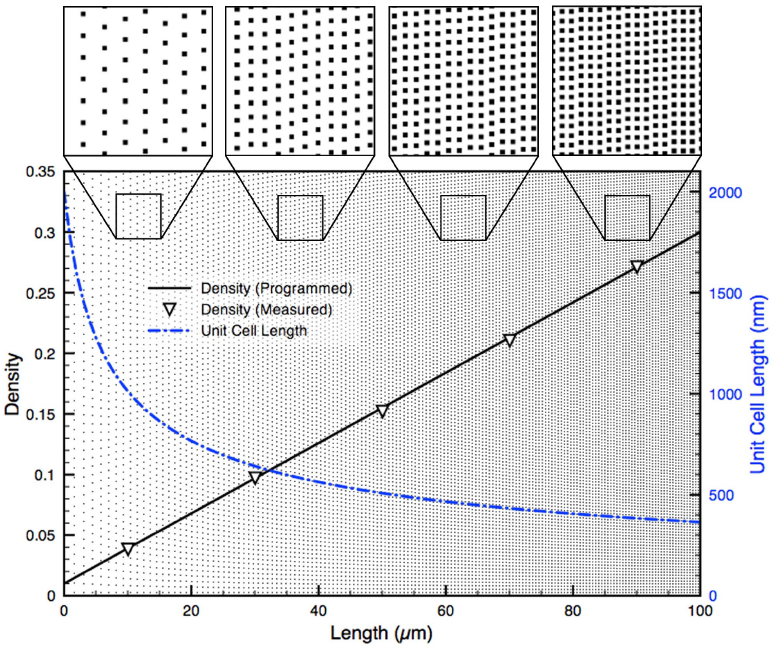
Programmed and measured density for an ordered linear DNG. A 100 × 100 µm^2^ *linear DNG comprised of* 0.04 µm^2^ nanodots and spanning from 0.01 to 0.30 density. The measured density (left axis) precisely matches the programmed density (R^2^=0.9998). The unit cell dimensions (right axis) change with the inverse square root of the density. Top insets show various spacing of nanodots at positions along the DNG.

To verify that ordered gradients matched the programmed density functions, gradients were exported as bitmap images and the ratio of black to white pixels, averaged over several columns of nanodots, was used to measure density. The resulting gradients were found to match the programmed density functions with high fidelity (*R^2^=0.9998*). Using this algorithm, a density of zero cannot be reached, as it would require an infinitely large unit cell. Consequently, when designing gradients of fixed length, the lowest density that is accessible may be limited since large unit cells are required at very low density.

### Random Gradient Algorithm

Naturally occurring gradients appear continuous at the microscale, but are in fact digital at the scale of molecules and proteins. The diffusion of biomolecules is subject to Brownian motion, and is therefore expected to be random. In an effort to mimic the nanoscale noise present *in vivo*, we present an approach to produce DNGs with fully randomized nanodot positions. Similar to ordered DNGs, random gradients have increasing density along the length and constant density along the width.

Several strategies were considered to produce randomized DNGs. The first was akin to the one proposed by Dalby, *et al.* [16] and consisted of starting with a regular distribution of dots, and randomly displacing the nanodots within their “unit cell” so as to avoid overlap with neighboring dots. However because this approach only produces a quasi-random distribution, it was not pursued. A second approach that was considered was to start with a pseudo-random dot pattern, and then compensate for the overlap by locally moving dots. This parallels a strategy used in back-lit displays [29], however these patterns are unsuitable for our application due to the dependence on or optimization for quasi-random distributions. Whereas we don’t see fundamental obstacles in developing an algorithm to randomly redistribute overlapping dots, such algorithms tend to be computationally intensive as the position of each dot needs to be tested against each other dot, and require multiple iterations to achieve the final distribution [27]. We thus devised a novel algorithm to create pseudo-random nanodot patterns for cell navigation that could translate arbitrary density functions into random nanodot patterns, and that was computationally economical.

The algorithm we developed compensates for the random overlap of dots by adding an excess of dots to achieve the programmed density. The algorithm first defines equally sized boxes to smoothly replicate the programmed density function without visible steps. The number of nanodots within each box was calculated based on the density function with predicted overlap at the box’s position, and random *x* and *y* coordinates were generated for each nanodot. Only the nanodot coordinates were stored. Noteworthy, the shape of the nanodots can be defined subsequently as the exported CIF file allows drawing shapes such as circles, polygons, etc. at the stored coordinates, thus making this approach flexible while reducing the file size and computation time. A consequence of the random nanodot position is that overlap between nanodots is possible. The frequency of overlap increases with density, which in turn results in a lower density than expected based on the surface area of all nanodots. However, because the distribution of the nanodots is random and the density known, the overlap is predictable, and can be compensated for, so that the designed DNG matches the density function with high fidelity. For each box with area *A_box_*, we first compute the probability *P_cov_* that any given point in the box is covered by a nanodot of area *A_dot_* (Eq. 3).

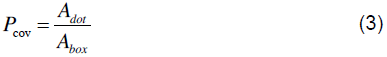

The probability *P_not cov_* that this point will *not* be covered by the nanodot is simply *1 – P_cov_* (Eq. 4).

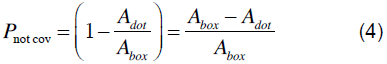

The probability that this point will not be covered by *N* nanodots simultaneously can then be found (Eq. 5).

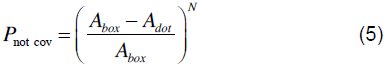

To determine the total area of the box covered by nanodots (*A_cov_*), the probability that a point *will* be covered is integrated over the area of the box (Eq. 6).

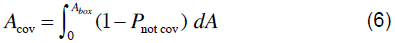

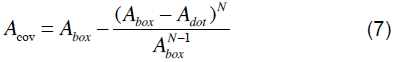

Eq. 7 can then be solved for *N* to determine the number of nanodots required for a given *A_cov_* in each box.

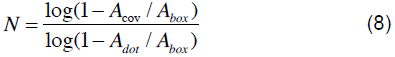

Using Eq. 8 it is possible to obtain the number of nanodots required for any given density by substituting *D* for *A_cov_/A_box_.* As the area occupied approaches the area of the box, the number of nanodots required increases rapidly, and is infinite for a density of 1. The maximum density for a DNG was thus set to 0.9999. Without compensation, at density of 0.9999 would require 9,999 nanodots of 200 × 200 nm^2^ to be seeded into a 400 × 1 µm^2^ box to reach this density. When accounting for the overlap, 92,099 nanodots are required, roughly a tenfold increase. Thus, producing a typical linear gradient, ranging in density from 0.0002 to 0.4444 requires 1,061,307 nanodots and a computation time of 4.56 min. The ordered algorithm for the same linear gradient requires 885,737 nanodots and a computation time of 3.88 min. Therefore, the random gradient algorithm overall required a 19.8% increase in nanodots and a 17.5% increase in computation time using our workstation. It is expected that the time will increase for random DNGs that reach a higher density.

We compared the programmed density with the measured density of a monotonic exponential curve with decay constant *k* (Eq. 9) using both the ordered and random gradient generation approaches. Eq. 9 is normalized to span the dynamic range over the length of the gradient. The random gradients produced with this algorithm were found to follow the programmed curve with high fidelity (*R^2^ = 0.99986*), and accurately match the measured density of ordered gradients produced with the same input function (Fig. 3).

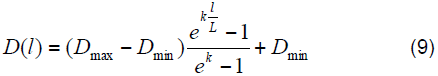

**Figure 3:**
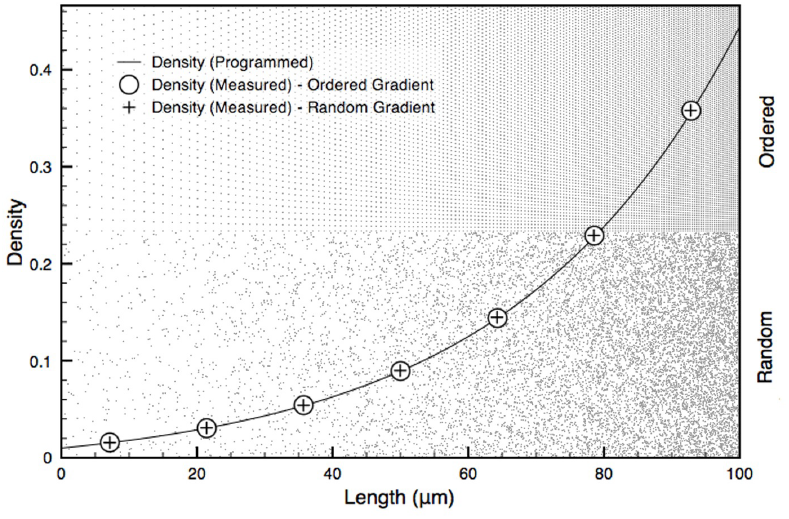
*Ordered and random DNGs superposed with their programmed exponential density function and actual, measured density.* A 100 × 100 µm^2^ exponential (k=3) DNG spanning from a density of 0.01 to 0.30 *is shown in ordered (top half) and random (bottom half) form. The random DNG was programmed by subdividing it into 1 µm wide boxes with random seeding of nanodots and compensating for overlap*. The measured density follows the programmed exponential curve with high fidelity for both ordered (R^2^=0.99996) and random (R^2^=0.99986) DNGs.

The distribution of neighboring nanodots can be parsed to verify whether it satisfies the conditions of randomness. Since density changes along the length of the DNG, randomness can only be assessed in the direction of constant density, perpendicular to the gradient. The randomness along each box was verified using Ripley’s K function based on the number of nanodots *N_pi_* within a distance *s* from the *i^th^* point *p_i_* taken over the sum of all points *n* and normalized by the area *λ* (Eq. 10).

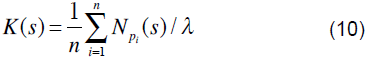

For a homogenous, random Poisson distribution, the expected value of Eq. 10 is *πs^2^*. Deviations from *πs^2^* indicate regions of clustering or dispersion [31]. Using the coordinates of nanodots from the DNGs, *K(s)* was found to lie within a 95% confidence interval obtained from 10 simulations of randomly distributed coordinates at the same density. This suggests the nanodots are spatially homogenous and randomly distributed compared to ordered gradients, which lie outside the confidence interval (Fig. 4).

**Figure 4:**
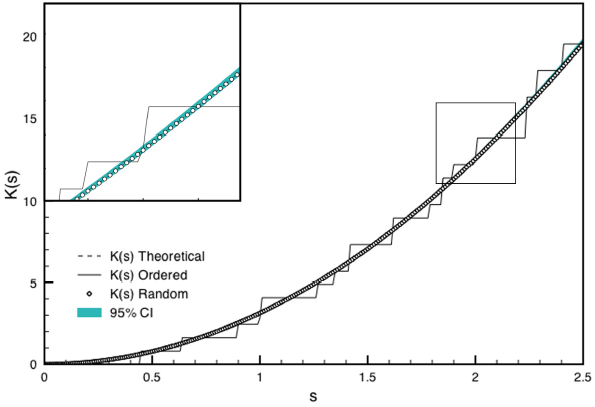
Ripley’s K analysis for randomness and spatial homogeneity of a sample box produced with the ordered and random DNG algorithms. K(s) for ordered and random gradients for a 10 × 400 µm^2^ box at 0.20 density are shown. A 95% confidence interval was calculated as 1.96 times the standard deviation of K(s) from 10 simulations of randomly distributed points at 0.20 density. The inset shows a magnified portion of the graph indicated by the box.

**Figure 5:**
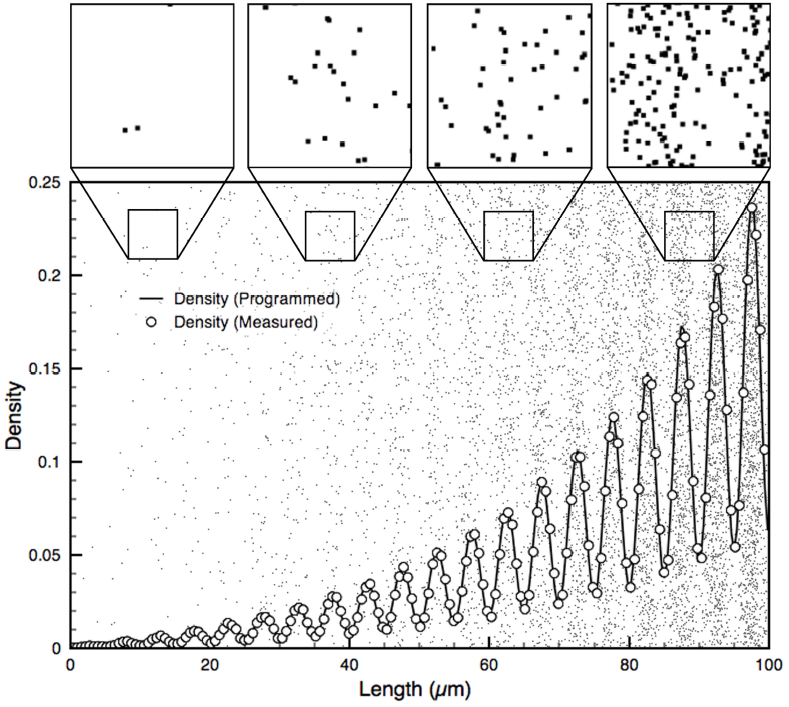
Non-monotonic random DNG superposed with its density function and actual, measured density. A 100 × 100 µm^2^ non-monotonic DNG. The programmed density function (line) is a sinusoid (A=0.10, B=20) with an average exponential trend (k_1_=3) and exponentially increasing amplitude (k_2_=3). Measured density from bitmap (circles) accurately follows the programmed function (R^2^=0.9988). Insets show close-up views of nanodots and reveal increasing overlap of nanodots at higher densities.

### Non-monotonic gradients

We have shown that linear and exponential gradients can be produced using either the ordered or random algorithms. These curves are monotonic, meaning they only ever increase or decrease. Given the approaches outlined here, more complex gradients can be easily generated from any input density function, specifically non-monotonous curves. To study the ability of cells to recognize an average gradient in a non-monotonous environment, we propose linear and exponential gradients that are superposed with sinusoidal functions with either constant amplitude, or with linear or exponentially changing amplitude. In the anticipation of future cell experiments, we also programmed a series of sinusoids with varying frequency and amplitude, while having an average slope of zero to act as negative controls.

Here, we demonstrate the flexibility of the algorithms with the production of non-monotonic gradients. One such gradient produced is a sinusoid with exponentially increasing amplitude superposed an exponential gradient. Eq. 11 gives the input density function for such a curve, where *A* is the amplitude, *B* is the number of oscillations, and *k_1_* and *k_2_* are the decay constants for the average gradient and amplitude of the sinusoid, respectively.

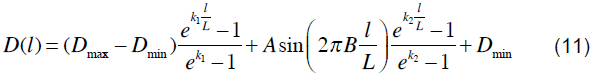

### One-Hundred-Gradient Array

The flexibility of the gradient algorithms and the fabrication method discussed below was leveraged by producing an array of 100 distinct gradients within a 35 mm^2^ area (Fig. 6, Table. S1). The array is comprised of 20 ordered and 80 random gradients, with densities ranging from 0.0002 to 0.4444 for a maximum dynamic range of 3.85 OM. For ordered gradients, this corresponds to a maximum pitch of 14.8 µm at low density, and 100 nm at high density. The minimum density is limited by the size of the cell and its capacity to sense the gradient, *i.e.* the cell will not sense the gradient if dot spacing exceeds one third the size of the cell, which we previously assessed using the migratory response of C2C12 myoblasts [14]. The maximum density is defined by the resolution of e-beam lithography, which limits the minimum pitch to 100 nm. To address how randomness affects cell navigation, 20 gradients (10 linear and 10 exponential) were produced as both ordered (#1-20) and random (#81-100) gradients. This portion of the array will serve to address gradient sensing mechanics for cell migration that may arise from either (i) the absolute concentration of the gradient at a given location or (ii) the concentration ratio between the cell’s leading and trailing edges.

**Figure 6:**
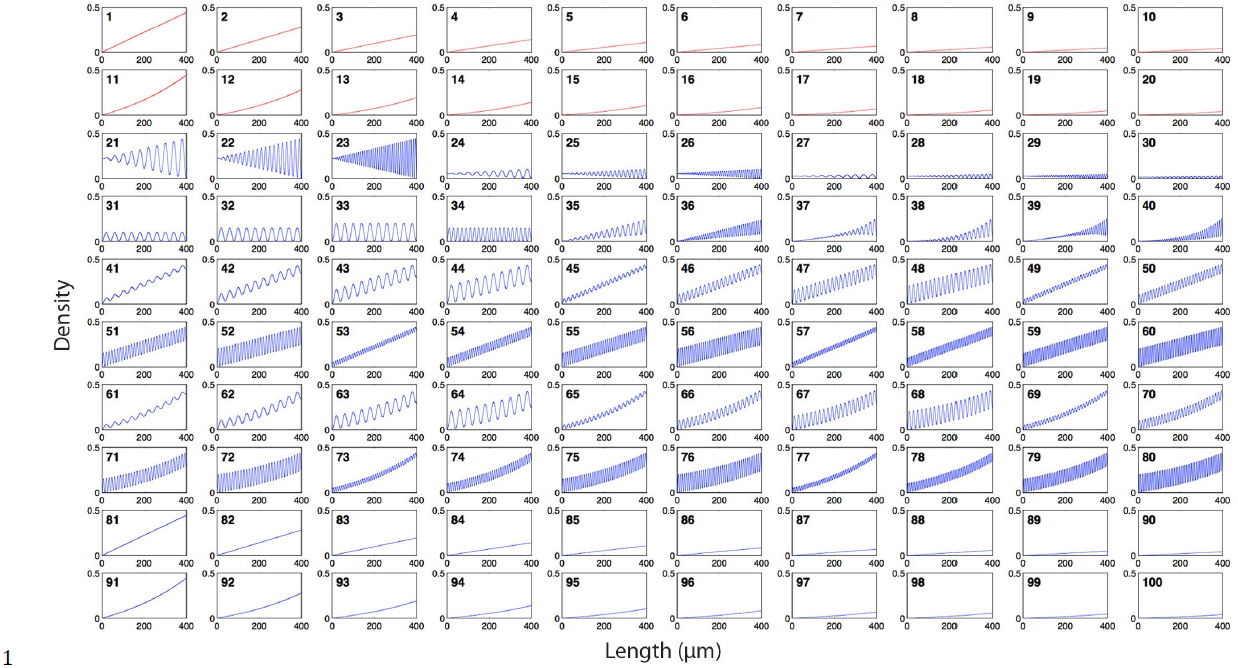
Density functions of the array of 100 DNGs. Each box shows the density function of one gradient. Functions 1-20 were produced with the ordered gradient algorithm (red), and functions 21-100 were produced with the random gradient algorithm (blue). Functions 1-10 and 81-90 are linear. Functions 11-20 and 91-100 are exponential. Functions 21-34 are sinusoidal with no slope (controls), where 21-30 have linearly increasing amplitude and 31-34 have constant amplitude. Functions 35-36 feature a linear slope superposed with a sinusoid that has linearly increasing amplitude. Functions 37-40 feature an exponential slope superposed with a sinusoid that has exponentially increasing amplitude. Functions 41-60 have a constant slope superposed with a sinusoid of constant amplitude. Functions 61-80 are exponential slopes superposed with sinusoids of constant amplitude.

The other 60 gradients are non-monotonic and random. These consist of 14 sinusoids with no average slope (#21-34) that serve as controls. These vary in frequency and amplitude, and may have constant or increasing amplitude along their length. Controls for ordered gradients were designed in our prior work. The remaining 46 gradients (#35-80) have sinusoidal curves with various levels of complexity. These include sinusoids with linearly (#41-60) and exponentially (#61-80) increasing average density, and non-monotonic functions with linearly increasing amplitude and average density (#35-36) and exponentially increasing amplitude and average density (#37-40) to demonstrate the flexibility of this new approach in gradient generation. As discussed, the sinusoidal curves with different amplitude and frequency may introduce obstacles for cells trying to sense the overarching gradient, and replicate effects that may occur due to cell “mosaicism” in a controlled manner.

### Gradient Array Fabrication

The hundred-gradient array was etched 100 nm deep into a Si wafer by e-beam lithography (Fig. 7). The integrity of individual dots for ordered and random gradients was confirmed by scanning electron microscopy (SEM) (Fig. 8).

**Figure 7:**
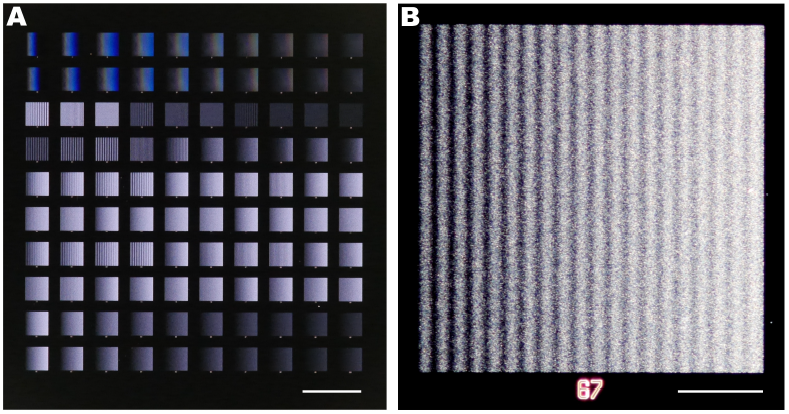
Optical images of the 100-gradient-array. The 100 DNGs were patterned into a Si wafer using e-beam lithography. (A) Image of all gradients; scale bar is 1 mm. (B) Dark-field image of DNG 67 which is a random sinusoid with exponentially increasing average density; scale bar is 100 µm.

**Figure 8:**
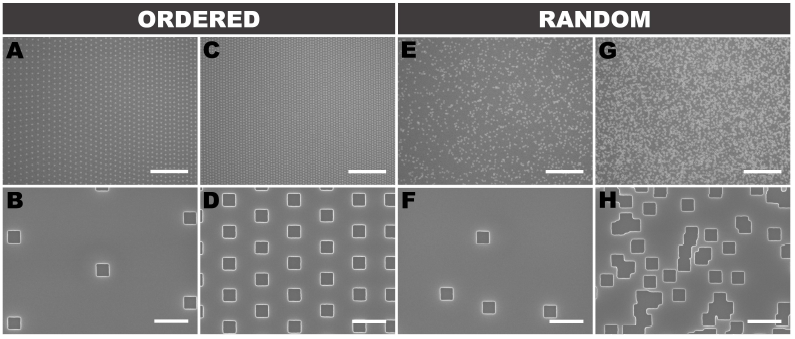
Nanodot distribution in ordered and random DNGs. SEMs of ordered (A-D) and random (E-H) exponential gradients at low (A,B,E,F) and high (C,D,G,H) densities. (H) highlights the random overlap between nanodots that is compensated for by the random DNG algorithm. Scale bars are 10 um (top row) and 500 nm (bottom row).

To translate the etched Si wafer into substrate-bound protein gradients, we employed lift-off nanocontact printing [14]. First, a PDMS intermediate replica was produced, followed by a second replication into low-cost Norland Optical Adhesive. A flat PDMS stamp was inked with a protein solution, and through contact with the plasma-activated NOA lift-off stamp, a monolayer of protein was selectively removed from the surface of the flat stamp leaving behind the digital nanodot protein pattern. Next, the flat PDMS stamp was brought into contact with a plasma-activated glass slide to transfer the DNG pattern. To confirm the accuracy of the replication and printing process, images of the design and the printed protein DNG were digitally overlapped and compared (Fig. 9), revealing a high fidelity between the two patterns.

**Figure 9:**
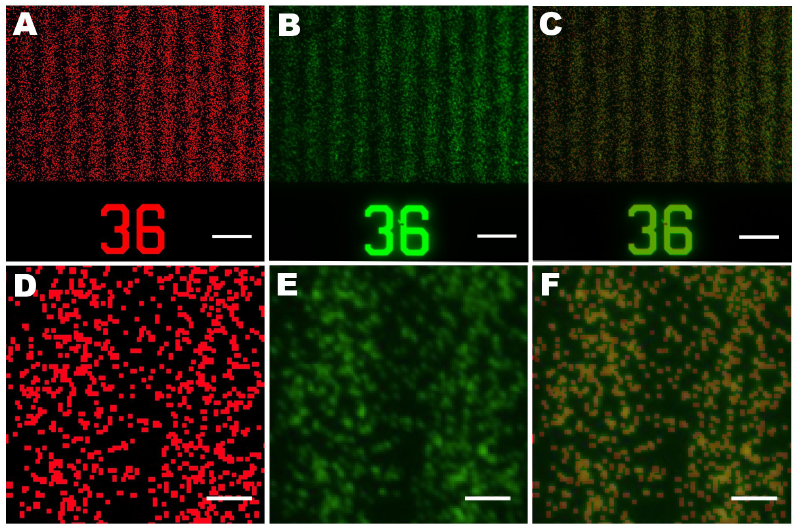
The printed nanodot pattern accurately replicates the design. Bitmap image of the programmed design colored in red (A, D) compared with a fluorescent image of nanocontact printed IgG by lift-off (B, E, green) and merged (C, F). In the inset (E), out of ∼1000 spots, 21 are missing, indicating that the replication works well. Scale bars A-C are 10 µm, scale bars D-F are 2 µm.

The pattern overlay in Fig. 9 was characterized by thresholding the printed image with boundaries of 31 and 255 in ImageJ and comparing it to the bitmap image (Fig. S3). The total number of nanodots is difficult to determine owing to the random overlap, but we estimate that there are ∼1000 spots, and using the above threshold, 21 were not found in the print. Overall, the transfer process is thus accurate to ∼98% in this example. Dust particles on the Si wafer or on the intermediate PDMS replica, or mechanical damage due to the replication process could account for the absence of these protein dots. Whereas the fidelity of the replication and printing process is thus high, it might be improved further by employing more durable polymers during the replication process as well as by working in a cleanroom environment throughout.

## Conclusion

Patterned substrate-bound protein gradients are a valuable tool to study a number of biological processes such as neuronal development or regeneration. The novel algorithms presented here, providing pseudo-randomness at a scale commensurate to the cellular level, can provide control over geometry and noise in DNGs. Combined with high-throughput patterning, an array of one hundred DNGs with linear, exponential, and non-monotonic slopes featuring 57 million spots over an area of 35 mm^2^ can be patterned at once in a matter of minutes. The diversity of DNGs shown here will help study and quantify the mechanisms by which cells sense and navigate through immobilized gradients. There are many opportunities for refining digital nanodot patterns. Firstly, to mimic the self-repellent nature of proteins adsorbing to surfaces, it might be useful to develop an algorithm that prevents, or limits, the overlap of nanodots. Secondly, whereas here two sinusoidal curves were superposed in one direction, such curves might be generated in two perpendicular directions and create two-dimensional navigation landscape while better mimicking local accumulation of guidance cues as puncta. Thirdly, it might be possible to program density functions that introduce clusters of noise that more accurately replicate the noise and mosaicism imposed by individual cells *in vivo* which can extend over tens of micrometers. Fourthly, it should be possible to pattern overlapping DNGs of different proteins that run in the same, or different directions, as well as produce any type of navigational landscapes that are found *in vivo* simply by converting the recorded densities into digital nanodot patterns, following the trend of rapid prototyping of replicas of living tissues [32]. Before expanding the nanodot patterning, it will be important to test and validate the current DNGs and establish the optimal conditions along with the suitable non-patterned reference surfaces for each study [33].

## Acknowledgments

We acknowledge Nabila Zaman for her help with gradient algorithms, and Marta Garnelo-Abellanas and Veronique Laforte for proofreading this article.

## Supplementary Material

**Figure S1:**
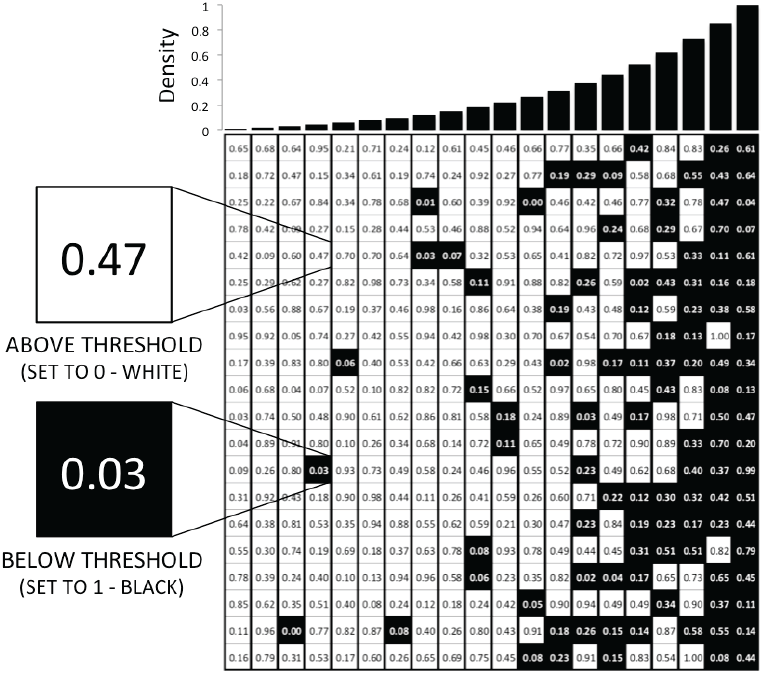
Random gradients produced by a random matrix threshold approach. A matrix of pseudo-random numbers is generated. Values greater than or equal to the density threshold are set to 0 (white), while values less than the threshold are set to 1 (black). The binary array can then be exported directly as a bitmap image file. Each nanodot is represented by one pixel. Thus, for a 400 × 400 µm^2^ sized area with 200 × 200 nm^2^ nanodots, a 2000 × 2000 matrix with 4 million values is required. This approach does not provide a fully random configuration since nanodots are aligned to a grid. While these patterns appear random to the eye, the underlying grid might be sensed at the cellular scale. Theoretically, it is possible to further randomize the position of dots by subdividing the area, e.g. using a 50 nm grid to position and draw 200 × 200 nm^2^ pixels, but this would come at the cost of increased computational requirements and the possibility of overlap between adjacent dots.

**Figure S2:**
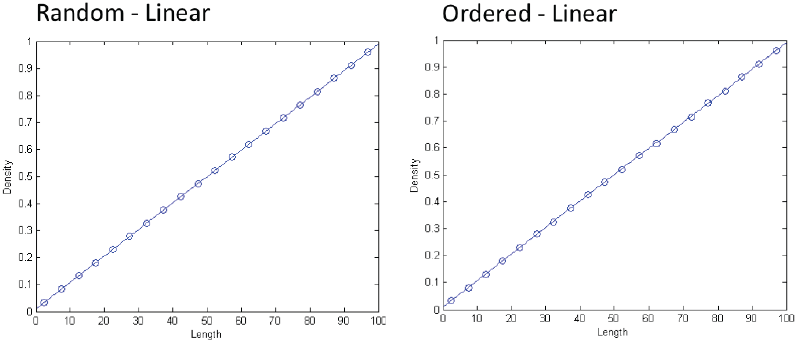
Ordered and random measured density matches programmed density over full range. Linear gradients from densities of 0.01 to 0.99 were shown to match (dots) the programed functions (line) with high fidelity for both ordered (R^2^ = 0.9988) and random (R^2^ = 1.0000) gradients. Using either approach, a high dynamic range can be achieved with near perfect match to the programmed function.

**Figure S3:**
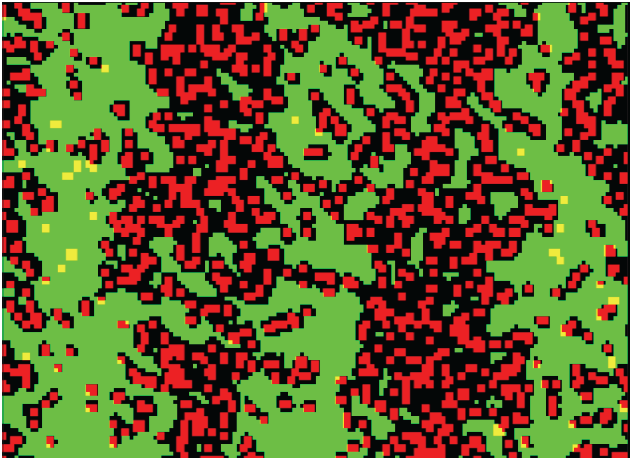
Image processing procedure to assess alignment of the DNG design and print. The fluorescent image of nanocontact printed IgG was first thresholded in ImageJ with boundary values of 31 and 255. The image was then transformed to binary and the binary values inverted to facilitate visualization. The edited fluorescent image (green) was then merged with the bitmap (red) and yellow dots, indicative of non-printed dots, were counted to determine to what extent the print matched the design.

